# Eelgrass seeds host a distinct microbiome that is consistent along a salinity gradient

**DOI:** 10.64898/2026.01.19.700372

**Authors:** Anne Brauer, Katharina Kesy, Mia M. Bengtsson

**Author notes:** **Author contributions** AB conceived the study together with MMB, carried out fieldwork, laboratory- and data analysis and wrote the manuscript. KK contributed to data analysis and interpretation. MMB supported experimental work and data interpretation and wrote the manuscript. All authors edited the manuscript and approved the final version.

## Abstract

Seagrasses are the only plants that flower and produce seeds in the marine environment, where they form vast meadows that fulfill important ecosystem functions. Yet, seagrass cover has declined in many coastal areas around the world and re-colonization is slow. Despite clonal growth, seed recruitment is essential for seagrass dispersal and regional genetic diversity. While the seed microbiome of several terrestrial plants has been shown to influence germination and seedling survival, the role of the seagrass seed microbiome is still unclear. We investigated the microbiome of eelgrass (*Zostera marina*) seeds, leaves and roots along the natural salinity gradient of the German North and Baltic Sea coasts. Despite this strong variability, *Z. marina* seeds harbor distinct prokaryotic and eukaryotic microbial communities compared to those on leaves and roots. Predicted microbial functions suggest roles in nutrient cycling and germination, which may be critical for recovery and restoration of seagrass ecosystems.

**Scientific significance statement:** Seagrass meadows are important but declining coastal ecosystems. Around the world, efforts are being made to restore seagrass meadows in order to halt the loss of biodiversity and maintain ecosystem function. With an evolutionary history on land, seagrasses have introduced unique features to the marine environment, such as seeds. In terrestrial plants, the seed microbiome has been shown to be important for seed germination and seedling health. Here we show that the northern hemisphere seagrass *Zostera marina* has a distinct seed microbiome, containing core bacterial and eukaryotic taxa and predicted functions that may play a role in seed germination and vertical microbiome transmission. Our results lay the foundation for future seagrass restoration efforts that seek to manipulate seed microbiomes to improve seedling survival.

**Data availability statement:** Raw DNA sequence data has been deposited in the European Nucleotide Archive (ENA) at EMBL-EBI under accession number PRJEB72211. Processed data and associated metadata will be made available on PANGEA via the German Federation for Biological data (GFBio.org).

## INTRODUCTION

Seagrasses make up a paraphyletic group of flowering plants that evolved to recolonize marine habitats during the last 100 mio. years (Larkum et al. 2006; Olsen et al. 2016). In that time, they have transformed coastal regions as ecosystem engineers providing unique habitats - seagrass meadows - that are marine biodiversity hotspots and offer numerous ecosystem services, including carbon sequestration (Hemminga and Duarte 2000; Duarte et al. 2005). The success of seagrasses can be explained, in part, by their adaptation of terrestrial evolutionary innovations such as seeds and roots to the marine environment. Seed-based reproduction allows them to maintain genetic diversity, facilitates dispersal and thereby, in combination with clonal growth, enables colonization of new areas. Nevertheless, seeds are also a vulnerable life cycle stage in seagrasses, as seed predation (Infantes et al. 2016), unsuitable environmental conditions (Jørgensen et al. 2019), and pathogen attack (Govers et al. 2016), for example, compromise seed germination and seedling survival. Despite many similarities to the seeds of terrestrial plants, seagrass seeds face unique challenges in the interaction with the coastal marine environment, which features steep gradients in salinity, inorganic nutrients and oxygen availability. Notably, they are subject to permanently hydrated conditions which offer a beneficial environment for microbial colonization compared to for terrestrial seeds which are at least periodically desiccated.

Like all other larger organisms, seagrasses live in close association with a community of microorganisms, their microbiota (Ugarelli et al. 2017; Tarquinio et al. 2019). Several beneficial interactions between microbial partners and seagrasses have been demonstrated, such as nutrient supply to seagrass roots (Mohr et al. 2021) and leaves (Pfister et al. 2023) and detoxification of sediments via sulfur oxidation (Jensen et al. 2007; Wang et al. 2021). Overall, the microbiome of seagrass vegetative tissues (e.g leaves, rhizomes and roots) reflects its marine environment (Fahimipour et al. 2017), and shares features with the microbiomes of macroalgae (Egan et al. 2013; Tarquinio et al. 2019), including abundant marine Bacteriodota bacteria (e.g. Chen et al. 2022), and a functional capacity of complex carbohydrate degradation (Crump et al. 2018; Miranda et al. 2022; Lu et al. 2023). However, in some instances, seagrass host-microbe interactions rather resemble those of terrestrial plants, such as the nitrogen-fixing root symbionts of the seagrass *Posidonia oceanica*, which are functionally analogous to terrestrial legume-rhizobia associations (Mohr et al. 2021). Like roots, seagrass seeds do not have a counterpart among macroalgae and there is currently little known about the similarities or differences of seagrass seeds to terrestrial plants. However, the seeds of the seagrass *Halophila ovalis* have been demonstrated to contain an endophytic microbiome, with several bacterial taxa implicated to provide beneficial functions for the host (Tarquinio et al. 2021).

Eelgrass (*Zostera marina*) is widespread across temperate seas of the Northern hemisphere, yet eelgrass meadow ecosystems are in decline, threatened by human activities and climate change (Orth et al. 2006; Dunic et al. 2021). Eelgrass meadow restoration using seeds may be a way to counteract biodiversity loss and restore ecosystem function, yet several technical challenges and low germination rates of seeds currently hamper these efforts (Marion and Orth 2010; 2012; Infantes et al. 2016). A better understanding of interactions within the seagrass microbiome has recently been suggested as a key factor in improving seed-based restoration success (Unsworth et al. 2023). However, we currently do not know how the eelgrass seed microbiome is structured, whether it is stable among populations and along environmental gradients and what role it may play in germination and seedling survival.

Here, we investigated the prokaryotic and eukaryotic microbial communities of eelgrass seeds in comparison to leaves and roots from different sites along the German coastline which features a natural salinity gradient. We aimed to identify core microbial taxa and predicted functions which define the eelgrass seed microbiota, regardless of the local environmental conditions. Specifically, we hypothesized that 1) seed microbiota would be more similar in composition to leaf than to root microbiota, because of their spatial proximity and shared environment in the water column. Further, we 2) expected to find previously detected potentially pathogenic taxa on seagrass seeds, but that these taxa would occur sporadically. Finally, we hypothesized 3) that bacterial functional traits of seed core microbiota would include potentially beneficial functions for germination and seedling survival, similar to those observed in terrestrial plants.

## MATERIALS AND METHODS

### Sample collection

We collected seagrass *Zostera marina* shoots at 10 sites along the coast of Germany (Fig 1, Table S1). Sampling was conducted during the flowering periods, which differ between the Baltic and North Sea, from June-July 2020 (Baltic Sea) and in September 2020 (North Sea). At each site we collected a reproductive and a neighboring vegetative shoot. From the reproductive shoot the spathe containing the seeds (referred to as sample type seeds or S) were sampled. From the vegetative shoot an up to 10 cm long tip from the youngest and a 10 cm tip of an older leaf (referred to as sample type young leaf YL and old leaf OL, respectively) as well as roots from 1 rhizome node near the meristem (referred to as sample type root or R) were cut off. Sample material was fixed in a high-salt preservation solution (6.16 M NH4SO4, 0.025 M EDTA, 0.025 M Na3C6H5O7). Fixed samples were brought to the laboratory and stored at 4 °C until DNA extraction.

**Figure 1:**
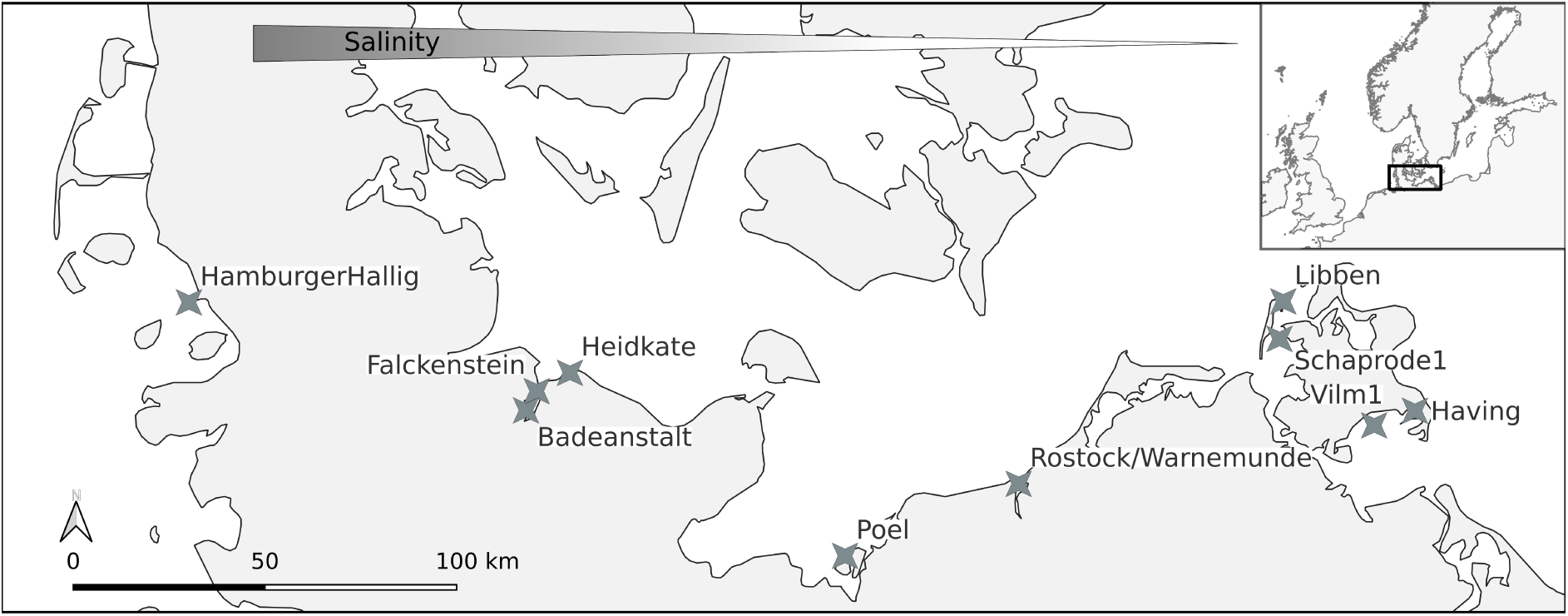
Map of the sampling sites in the southern Baltic Sea and North Sea (coastline of Germany), featuring a natural salinity gradient of 23 ppm at the westernmost sampling site Hamburger Hallig to 8.5 ppm in the East at Vilm1. Coordinates and measured environmental variables can be found in Table S1.

### DNA extraction and amplicon sequencing

For DNA extraction, seagrass leaves were cut into ∼2 cm long pieces, roots were detached from the rhizome and seeds taken out of the spathe. The samples were briefly soaked in PBS buffer (pH 7) prior to extraction, to wash off salts from the preservation solution. DNA was extracted using the DNeasy power soil pro kit (Qiagen) according to the manufacturer’s instructions, with minor modifications: To extract DNA of epibiotic microorganisms while avoiding coextracting seagrass DNA the samples were added to the extraction buffer, then sonicated on ice twice for 7 min and shaken at low intensity (FastPrep, MP Biomedicals, 30 s 4m/s). The plant material was subsequently removed from the extraction tubes before microbial cell disruption (45s at 5m/s). DNA was stored at - 80°C until sequencing. Amplicon sequencing of the V4 region of the *16S* rRNA genes and the V7 region of the *18S rRNA* genes (primer pairs 515f: 5‘-GTGYCAGCMGCCGCGGTAA-3‘, 806r: 5‘-GGACTACNVGGGTWTCTAAT-3’ (Walters et al. 2015) and F-1183mod: 5’-AATTTGACTCAACRCGGG-3’, R-1443mod: 5’-GRGCATCACAGACCTG-3’ (Ray et al. 2016), respectively) was carried out as described previously (Bengtsson et al. 2017, Brauer & Bengtsson 2022).

### Sequence analysis and functional prediction

Amplicon sequence reads were processed to identify unique amplicon sequence variants (ASVs) using the dada2 pipeline (Callahan et al. 2016, version 1.18) in R (R Core Team 2025). ASVs were taxonomically annotated in dada2 using the Silva database for 16S reads (Pruesse et al. 2007, silva nr99 v138.1, min Boot= 80) and the pr2 database for 18S reads (Guillou et al. 2013, version 4.13.0). 16S ASVs annotated as non-prokaryotic, including chloroplasts and mitochondria as well as 18S ASVs annotated as plants (*Embryophyceae*, seagrass host contamination) were removed before downstream analysis. Eukaryotic ASV’s of special interest without species assignment were compared against the core nucleotide (nr) database (Sayers et al. 2025) using BLASTN (NCBI, online tool, Camacho et al. 2009).

Functional pathway prediction was carried out using the Paprica pipeline (Bowmann and Ducklow 2015, Barbera et al. 2019, Czech et al. 2020, Guillou et al. 2013, Haft et al. 2018, Karp et al. 2021, Nawrocki and Eddy 2013). Paprica aims at inferring metabolic pathways by placement of ASVs in a phylogenetic tree of known bacterial genomes.

The number of ribosomal RNA operons per ASV was predicted using the rrnDB database v 5.8 (Stoddard et al. 2015). When available *rRNA* gene copy numbers at genus level were used, where genus level predictions were missing family level copy number was used. In the case that at both levels no database entry was found a *rRNA* gene copy number of 1 was assumed.

### Statistical analysis

All statistical analyses were carried out in R (R Core Team 2025). The vegan package (Oksanen et al. 2025) was used to calculate richness and evenness of the read based dataset, rarefied to even sampling depth of the smallest library. Bray curtis distance matrices on hellinger transformed relative abundance data were created for nmds (Non-metric multidimensional scaling) plots. Permanova was used to investigate the influence of sample type and salinity on the community composition. Pairwise permanova was calculated to check which sample types differ from each other using pairwiseAdonis (Martinez Arbizu 2017) with bonferroni correction. Differential abundance of AVS’s as well as of predicted functional pathways between the seeds and the other sample types were calculated with Maaslin2 (Mallick et al. 2021). To identify core and exclusive ASVs of the different sample types, venn diagrams were created using VennDiagramm (Chen 2022). Core ASV’s are referred to as ASV’s that occur on all samples of a specific sample type, exclusive ASV’s are the subset of core ASV’s that only occur on all samples of a specific sample type and are not core ASV’s of other sample types (but can be present in some samples of other sample types). Mean rRNA copy number of samples was calculated from predicted *rRNA* copy number per ASV multiplied with the rarefied abundance and tested with Anova and Tukeys-post hoc test.

## RESULTS

Out of 8312 total 16S ASVs (from here on referred to as “bacterial” as archaea only made up 0.6% of ASVs), 3915 occurred on seeds, and 200 only on seeds (Fig. 2a). We identified 16 bacterial ASVs which occurred in 100% of seed samples, referred to as “core” seed ASVs. Of these, 7 ASVs were present exclusively on all seed samples, and were defined as “exclusive core” seed ASVs (Fig. 2b). For eukaryotes, 358 out of 804 ASVs were detected on seeds, and 3 of these could be defined as core ASVs, of which 2 were also exclusive core ASVs. Seed bacterial and eukaryotic core ASVs are listed in Table 1.

**Figure 2:**
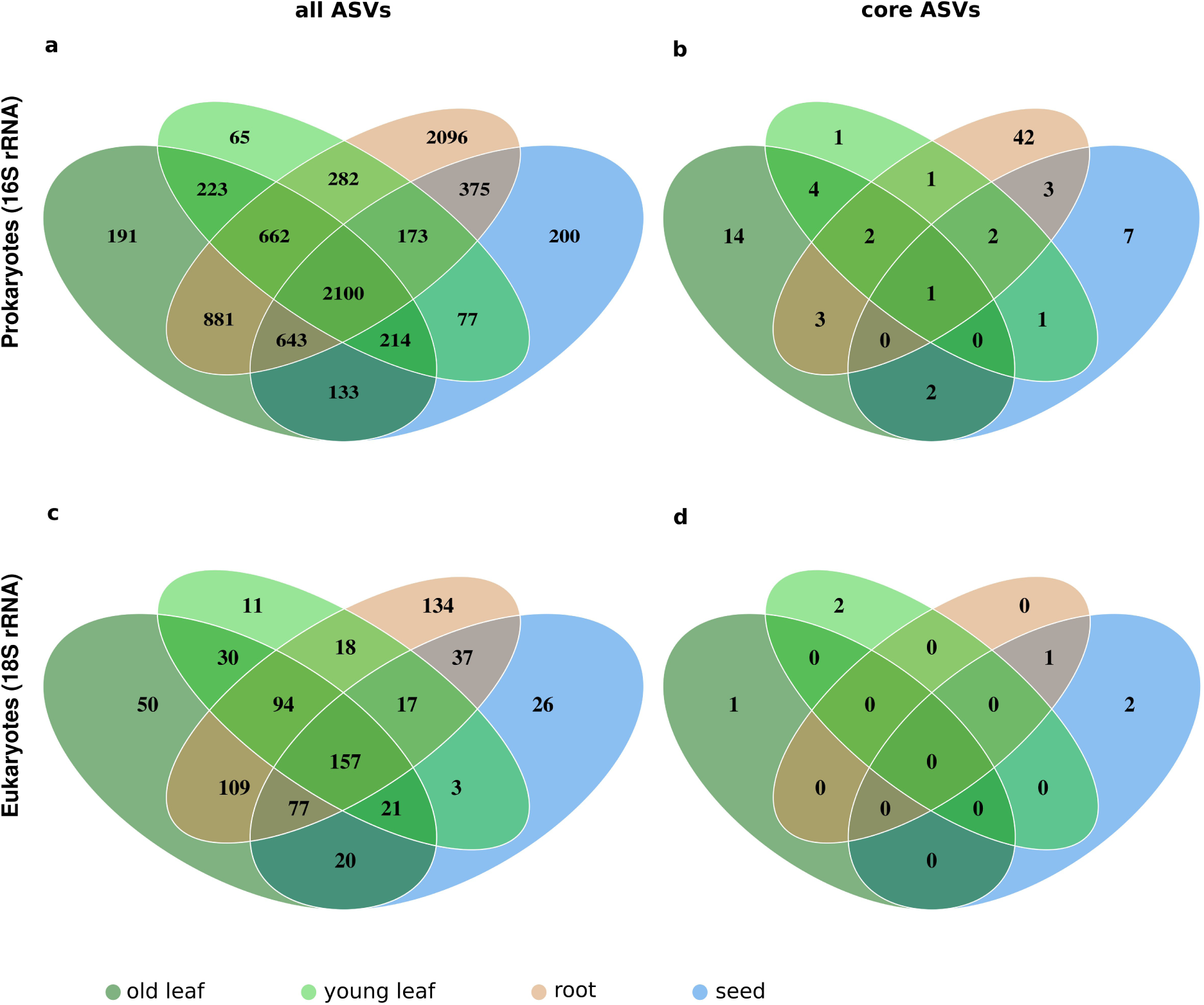
Number of all ASVs shared or exclusive for the different sample types based on bacterial (a) and eukaryotic (c) dataset and number of the ASVs that occur on all samples of either of the sample types shared or exclusive between the different sample types based on bacterial (b) and eukaryotic (d) dataset.

**Table 1:**
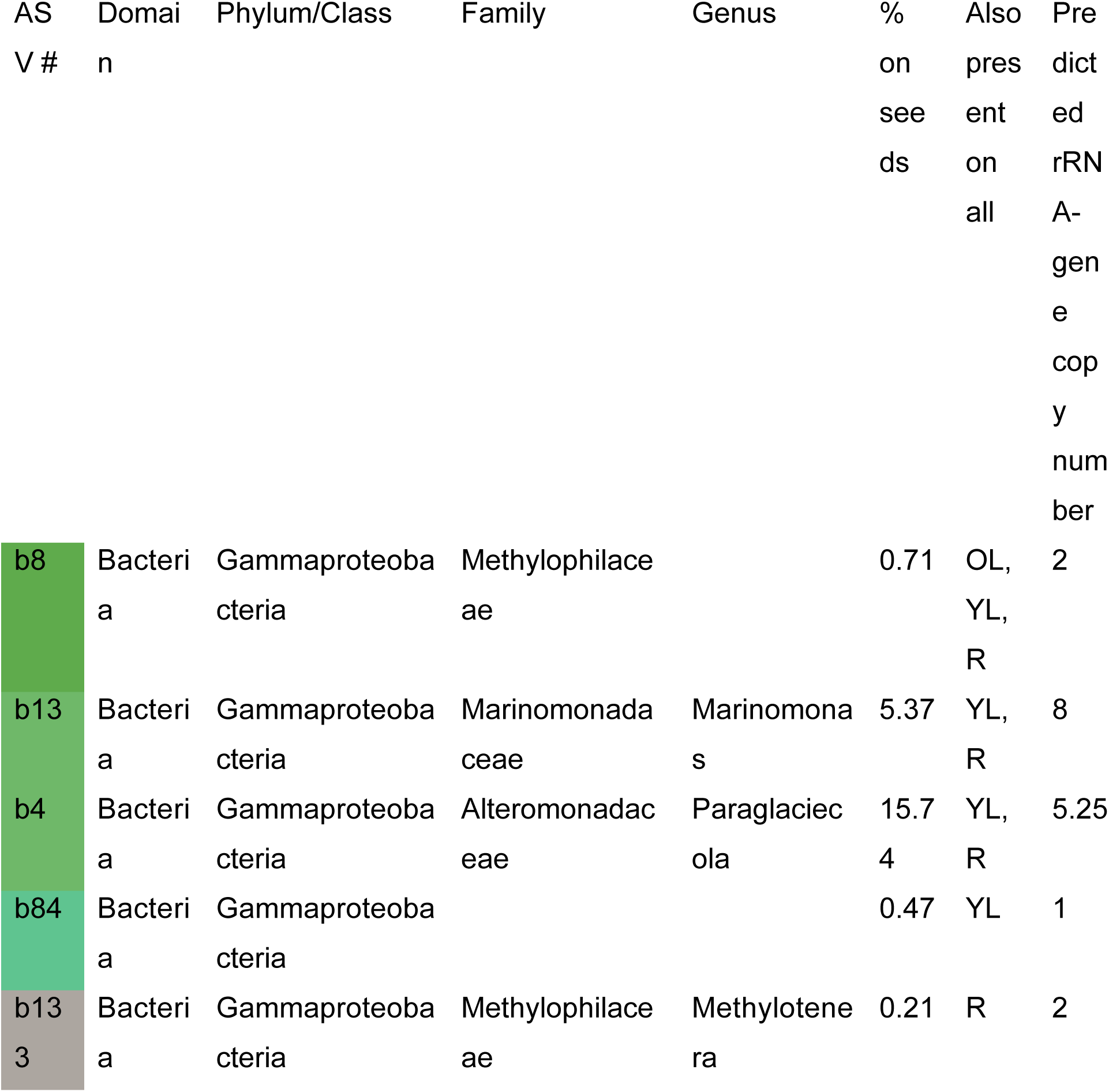
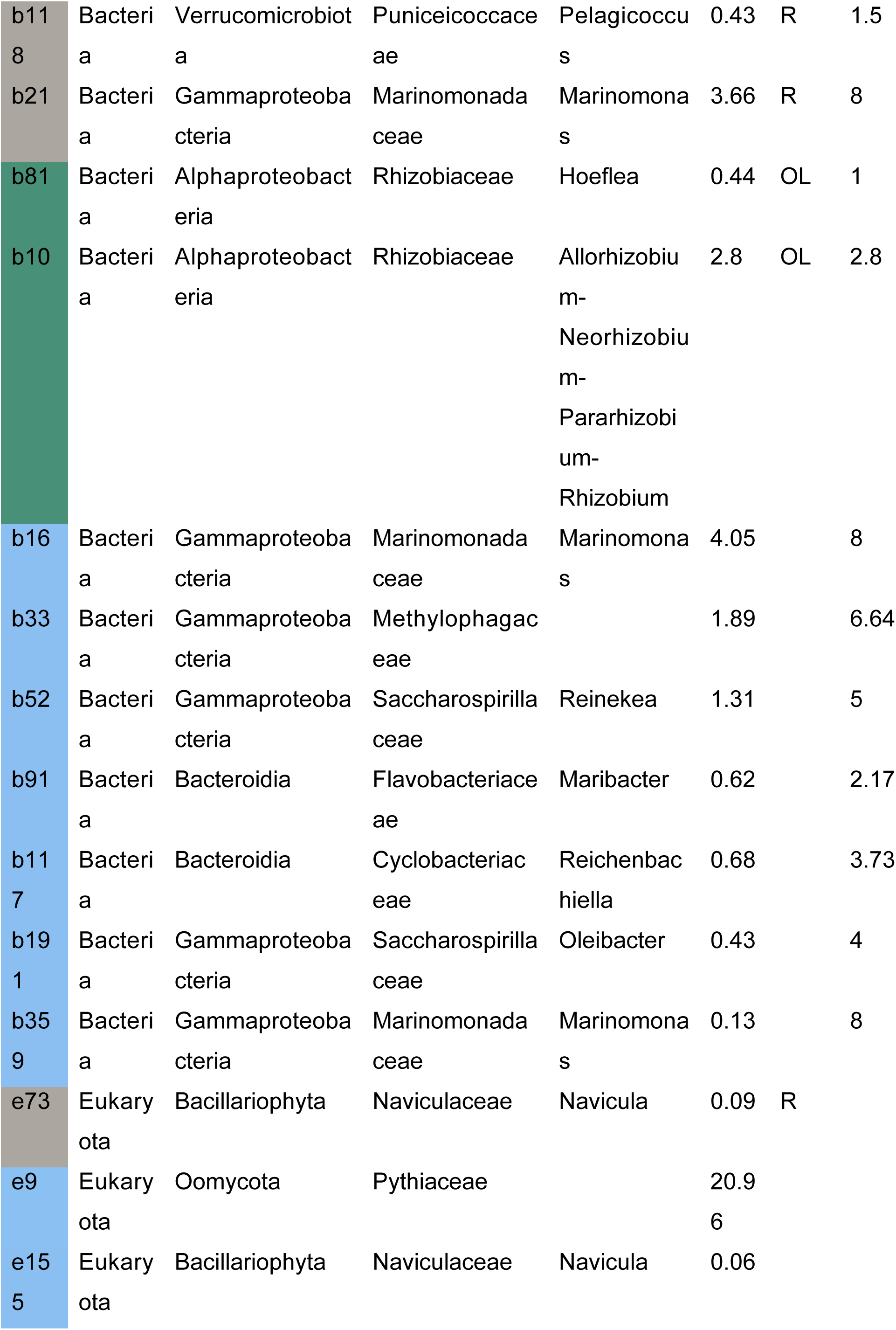
Taxonomic classification and relative abundance of core bacterial and eukaryotic ASVs on seagrass seeds (core defined as those occurring on 100% of the seed samples). “%” refers to the average relative abundance of the ASV in the seed samples, calculated separately for the 16S and 18S datasets. The colors in the first column correspond to the respective fields in the Venn diagram in Fig. 2 where blue indicates exclusive occurrence on all seeds (”exclusive core”). Column 7 indicates in which sample type the respective ASV occurred in all samples other than seeds OL – old leaf, R – root, YL – young leaf, S – seed.

Four bacterial ASVs belonging to the genus *Marinomonas* (b13, b21, b16, b359) were identified as core, and two of these were exclusive core ASVs. The most abundant core ASV was 100% identical to the type strain of *Paraglaciecola hydrolytica* (b4, 15.74% relative abundance on seeds), and was also detected on leaves (Table 1). Three core ASVs belonged to the family *Methylophilaceae*, and one of these (b8) was the only core ASV detected on all samples, regardless of sample type. Two core ASVs belonging to the *Rhizobiaceae* (b81, b10) were also detected on roots. Further core ASVs were classified as *Pelagicoccus* (b118), *Reineka* sp. (b52), *Maribacter* sp. (b91), *Reichenbachiella* sp. (b117) and *Oleibacter* sp. (b191). The most abundant eukaryotic exclusive core ASV was an oomycete of the family *Pythiaceae*(e9) (20.96 % relative abundance on all seeds). Further core ASVs comprised two diatoms of the genus *Navicula* (e155, e73).

Bacterial community composition varied mainly according to sample type, explaining 22% of variation (R2 = 0.22), while salinity explained 5% of variation according to PERMANOVA (p<0.01, Fig 3a). This pattern was similar for the eukaryotic communities (sample type 18%, salinity 5%, p<0.001, Fig. 3c). The seed microbiome was significantly different from old and young leaf, and root microbiome (pairwise PERMANOVA, p < 0.01 for prokaryotes and p < 0.05 for eukaryotes, Table S2 and Table S3, respectively).

**Figure 3:**
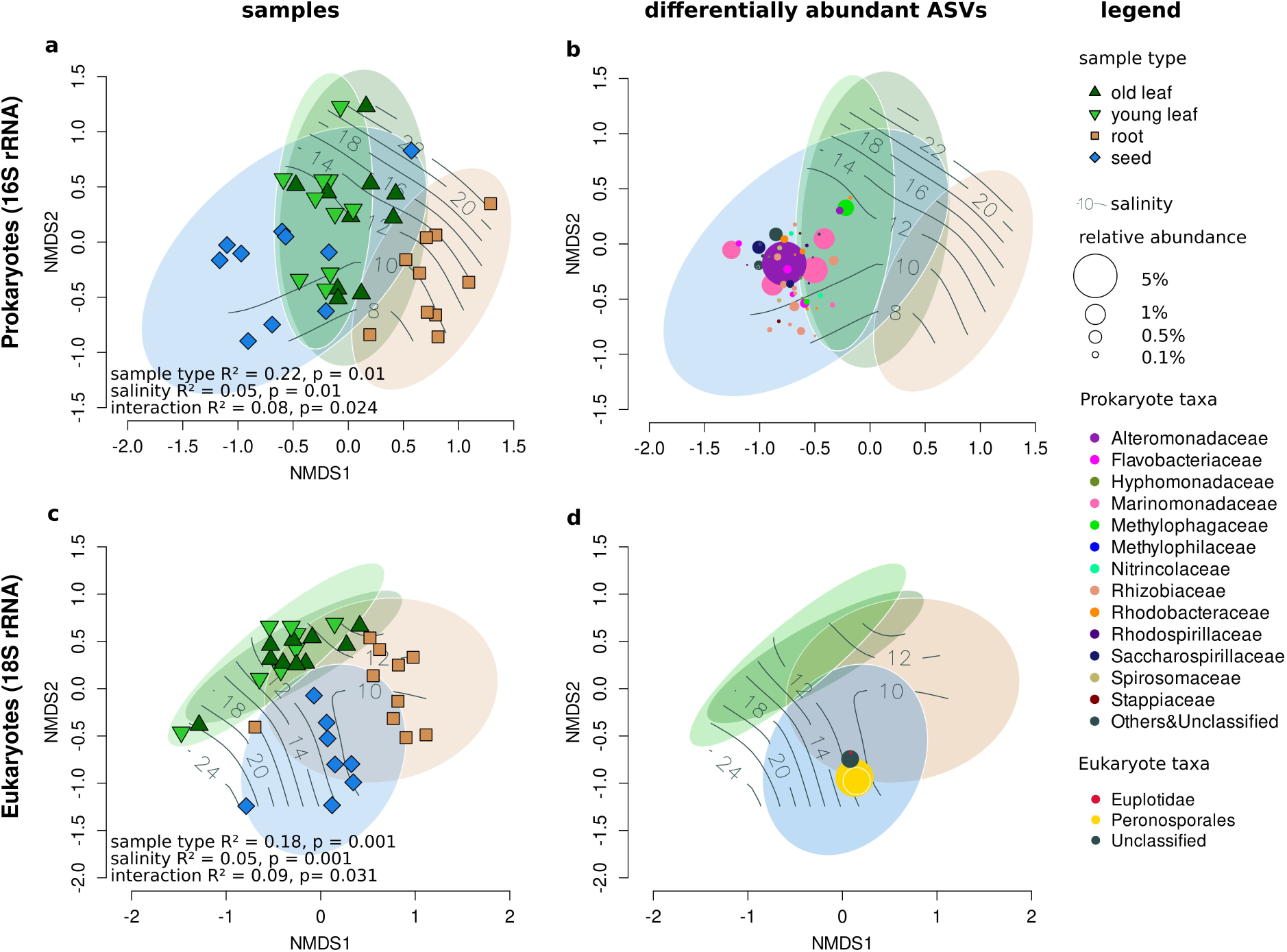
Variation in microbiome composition between sample types and along the salinity gradient is displayed in nMDS ordinations (bray-curtis dissimilarity) calculated based on bacterial ASVs (a-b) and eukaryotic ASVs (c-d). Ordinations represent dissimilarities between sites (a,c) and ASVs (b,d) that are differentially more abundant between seeds and all the other sample types. Point size corresponds to ASV overall abundance in the dataset, and color corresponds to ASV taxonomic classification. Ellipses indicate 95 % confidence intervals of site point position within sample types. Solid gray contour lines indicate fitted salinity values in ppm.

In addition to the core seed ASVs, the differential abundance analysis identified several bacterial ASVs which were significantly overrepresented on seeds, but were not necessarily part of the core microbiome (Fig. 3b, for a full list of differentially abundant bacterial ASVs see Table S4). Notable among these were Rhizobiaceae (11 ASVs), Rhodobacteraceae (6 ASVs), *Marinomonas* spp. (5 ASVs) and Methylophagaceae (2 ASVs). Only four eukaryotic ASVs were differentially more abundant on seeds compared to the other sample types (Fig. 3d, for a full list of differentially abundant eukaryotic ASVs see Table S5). Those were two oomycetes belonging to the family *Pythiaceae* (e9 and e18), a ciliophore of the genus *Euplotes* (e358), and an unclassified *Metazoa* (e51).

Functional pathway prediction revealed a distinct functional fingerprint of the seed microbiome (Fig. 4a), and reproduced the overall community composition patterns observed (Fig. 3a), with seed samples partially overlapping with leaf samples, and to a lesser extent with root samples. Functions corresponding to lipid and nucleotide biosynthesis, carbohydrate (e.g. alginate) and aromatic compound degradation and inorganic nutrient acquisition were more abundant on the seeds compared to the other plant parts (Fig 4a, for a full list of differentially abundant predicted functions see Table S6).

**Figure 4:**
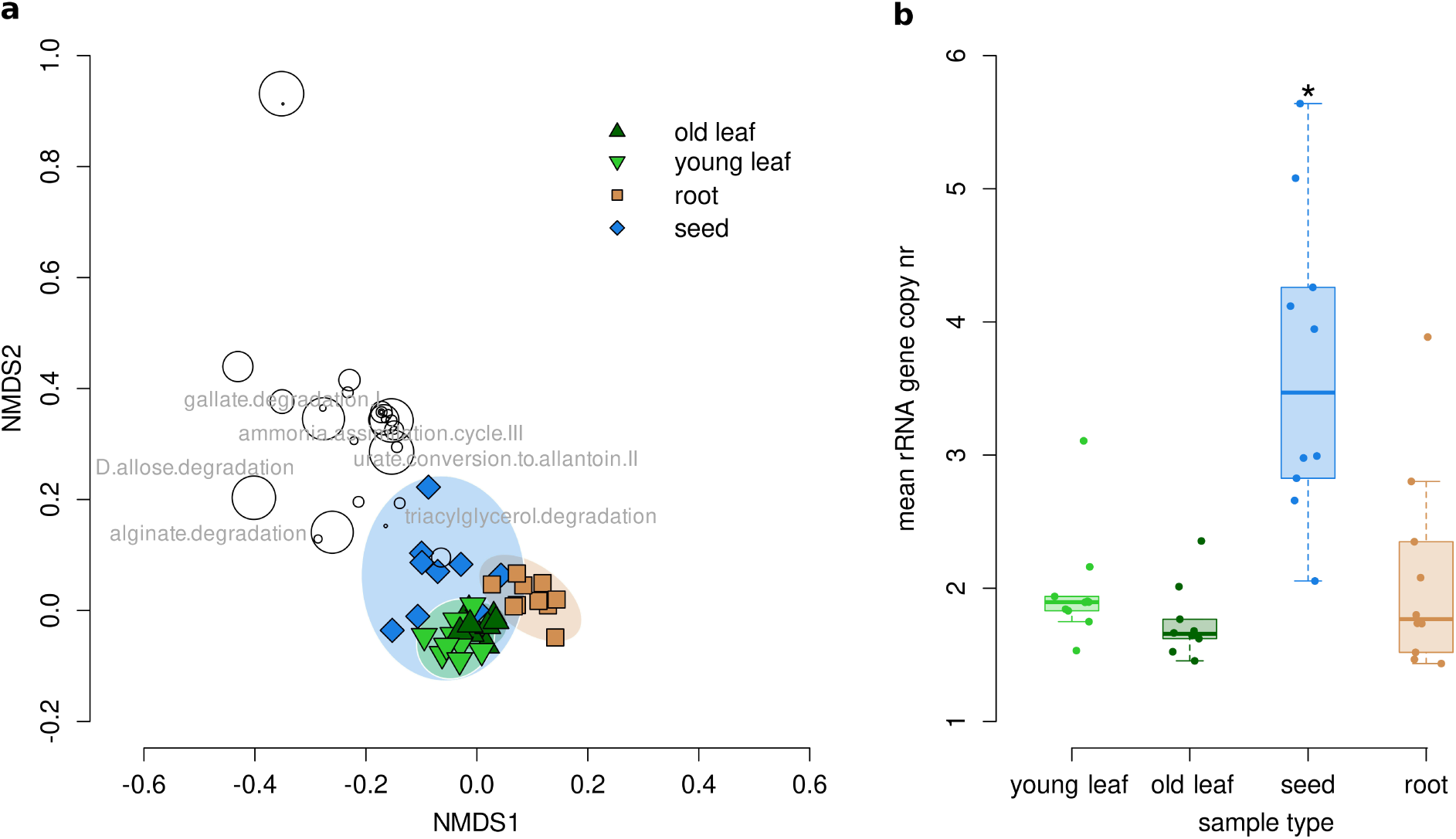
Variation in predicted functional pathways and rRNA operon copy numbers of bacterial ASVs. The nMDS ordination displays variation in functional pathway composition between sampling sites (squares and triangles) and predicted functional pathways (circles) which were differentially more abundant between seeds and all other sample types (a). Ellipses represent 95% confidence intervals of sample site point position. Size of the circles corresponds to the functions overall abundance. Selected functions are highlighted. Mean copy number of rRNA operons per sample (b).

Seed ASVs had a significantly higher mean number of predicted rRNA operons among all sample types (Anova p < 0.05, Fig. 4b).

## DISCUSSION

In this first investigation of the eukaryotic and prokaryotic microbial communities of the seeds of the seagrass *Zostera marina*, we found that the seeds have a consistenly distinct microbiota despite a strong environmental gradient (salinity). Our analyses showed that seed bacterial and eukaryotic microbiota composition was distinct from the leaf and the root microbiota. Contrary to our expectation that the seed microbiota would be similar to the leaf microbiota, there was not a strong overlap between leaf and seed microbiota composition, suggesting that the seeds are a unique microniche with different conditions than both roots and leaves. Despite an observed shift of the microbial communities with decreasing salinity we could identify a core microbiome consisting of taxa that appear on all seed samples.

The eukaryotic seed core microbiome was dominated by an oomycete ASV (ASV e9) from the family *Pythiaceae*, a genus that contains known plant pathogens (Thines and Kamoun 2010). Known putative seagrass pathogens include the oomycete genera *Phytophthora* and *Halophytophthora* (Govers et al. 2016) that can be combated by copper sulfate treatment during seed storage (Govers et al. 2017). We hypothesized that known seagrass pathogens would be detected on the seeds, but not be part of the core microbiome. As expected, members of the genus *Phytophthora* were detected on the seeds in varying abundances. More surprising is the high abundance of a *Pythiaceae* (oomycetes) core ASV on the seeds, raising the possibility thatall or most seeds seeds could be subject to infection. Oomycetes are widespread in terrestrial plant microbiomes (Durán et al. 2018; Sapp et al. 2018) and often considered pathogens (Thines and Kamoun 2010; Beakes et al. 2012) or saprotrophs (Hulvey et al. 2010). However, some members e.g. *Pythium* spp., can also be beneficial to plants by promoting plant growth and reducing biotic stress (Benhamou et al. 2012). With our methodological approach, we can not determine Which role the detected oomycetes play under the conditions present at sampling (see Judelson and Ah-Fong 2019). However, the role of these organisms within the seed microbiome needs re-evaluation, as for example, current Labyrintulomycetes are present, but not cause die offs in *Z. marina*, indicating a commensal role on seagrass leaves (Brakel et al. 2014). Likewise, members of the oomyctes prestent on seeds do not necessarily indicate infection, especially when being present in the core microbiome.

We expected bacterial seed microbiome members with predicted beneficial functional traits to be important on seeds, regardless of the salinity conditions. While our experimental approach using amplicon sequencing of the SSU rRNA does not allow direct observation of microbial functional traits, certain functions are taxonomically conserved or overrepresented within bacterial lineages, enabling prediction of traits based on literature and genetic information available through public databases.

Our findings highlight an important role of methylotrophic bacteria in association with seagrass seeds with two methylotrophic ASV’s in the seed core microbiota (b33 and b8). Methylotrophy appears to be an important functional trait in seagrass leaf microbiomes in general (Gebbe et al. 2025; Martin et al. 2020; Tarquinio et al. 2019; Crump et al. 2018; Bengtsson et al. 2017) where they likely consume methanol produced by the plant as a byproduct (Crump et al. 2018). Methylotrophs have also been identified as core members of the *H.ovalis* seed microbiota (Tarquinio et al. 2021). This is a shared feature with terrestrial plants where methylotrophic bacteria are omnipresent on seeds (Simonin et al. 2022) and can be transmitted from seeds to seedlings (Bajpai et al. 2025) where they can promote plant growth (Koshy et al. 2025).

Rhizobiaceae, which are well known for their interactions primarily within the rhizosphere of terrestrial plants (Spaink et al. 2012), including, but not limited to, nitrogen fixation (Díez-Méndez and Menéndez 2021) are other core members (b10 and b81) of the seed microbial community. Surprisingly, these ASVs were detected in root samples in much lower abundance, but were found highly abundant on seeds as well as old and young leaves. Rhizobia are also common core members on terrestrial plant seeds (Simonin et al. 2022) which can produce the plant hormone auxin (Ghosh et al. 2011) and fix Nitrogen (Díez-Méndez and Menéndez 2021).

Another prominent member of the core and enriched seed microbial communities are *Marinomonas* spp. (4 and 5 ASVs respectively). Several members of this genus are proven to have plant-growth promoting properties in seagrasses and coastal *Spartina* spp. grasses (Mesa et al. 2015; Lucena et al. 2016; Tsuchiya et al. 2024). *Marinomonas* spp. are also part of the core community of *H. ovalis* seeds (Tarquinio et al. 2021). *Marinomonas posidonica* improved the growth of *P. oceanica* seedlings (Celdrán et al. 2012), indicating that *Marinomonas* spp. are important microbiome members in several seagrass species.

A striking observation was the dominance of the seed microbial community by an ASV (b4) with 100% similarity to polysaccharide degrader *Paraglaciecola hydrolytica* whose type strain has been isolated from eelgrass leaves in Denmark (Bech et al. 2017). *P.hydrolytica* strain S66 was able to grow on several seaweed-derived polysaccharides (e.g. alginate, agar) as well as terrestrial plant polymers such as pectin. The presence of numerous genes linked to the degradation of these diverse and complex polysaccharides in its genome (which was included in the database used for functional prediction) may explain why the seed microbiota showed a higher predicted functional potential for degradation of some polysaccharides (e.g. alginate) compared to the microbiota of other plant parts in our study.

The higher average predicted *rRNA* gene operon copy number of the seed-compared to the other plant parts microbiota suggests that *Z. marina* seeds harbor microbial taxa with high potential growth rates (Roller et al. 2016; Klappenbach et al. 2000; Lauro et al. 2009). Fast growing bacteria with a copiotrophic lifestyle have been linked to germinating seeds of terrestrial plants (Torres-Cortés et al. 2018) with the number of fast-growing bacteria increasing during seedling emergence (Barret et al. 2015), reflecting the greater availability of labile carbon and nutrients compared to a dormant seed. In our study, the seeds were not in the process of germination, as they were harvested during maturation while still inside their spades. Unlike the seeds of most terrestrial plants, seagrass seeds do not undergo desiccation in their dormant stage, presumably offering an environment relatively rich in dissolved carbon and nutrients throughout maturation, dormancy and germination. This may pose a challenge to the survival of the seeds, making them exposed to opportunistic microbial pathogens which also feature high potential growth rates (Westrich et al. 2016). Seeds are indeed capsules of opportunity, which present colonizing microbes with two potentially attractive scenarios; Either the seed will germinate, exposing a rapidly developing seedling producing labile organic exudates as it starts photosynthesizing. Alternatively, the seed will die and be available to microbial degradation, releasing energy-rich stored organic compounds and nutrients. In either case, a high potential growth rate will allow associated bacteria to get a head start on capitalizing these unique resources in the marine environment.

Our findings offer strong indications that the eelgrass seed microbiome contains bacteria and microbial eukaryotes that interact closely with the seed and seedlings, thereby likely influencing germination, seedling survival and growth. We demonstrate that the seed microbiome is distinct from other plant parts and contain core taxa which are continuously present despite a strong environmental gradient. The seagrass seed microbiome bears functional resemblance to seed microbiomes of terrestrial plants, indicating that their shared evolutionary history has left an imprint in how the seagrass host interacts with microbes in its environment. This complements the understanding of the unique morphological and genomic adaptations of seagrasses that allowed them to colonize the marine environment (Olsen et al. 2016).

Identifying key microbial taxa and functions is the first step towards manipulating seagrass seed microbiomes to improve germination rates in seagrass restoration efforts. Microbial seed inoculation has been successfully used in terrestrial agricultural plants for productivity gains, including boosting germination rates (O’Callaghan 2016 and references therein). With typical germination success of only 1-5 % of sown eelgrass seeds (Orth et al. 2003; Infantes et al. 2016), there is great potential for improving seed-based restoration by increasing germination rates. Core seed microbiome taxa with potential beneficial functions, such as *Marinomonas* sp., would be preferred candidates for inoculation and germination trials with seagrass seeds. Targeted isolation of seagrass-associated bacterial strains in culture and screening for plant growth promoting properties should further help to identify strains suitable for seed inoculation in restoration trials. In addition, future investigations should look closer into the functional capacity and activity of natural seed microbiomes during the germination process using methods such as for example metagenomics, metatranscriptomics and metaproteomics to improve our understanding of the interactions between the microbiome and the seagrass host.

## Acknowledgements

The authors acknowledge the help of everybody who was involved in sampling, namely: Philipp Schubert, Laura Govers, Sven Dahlke, Philipp-Konrad Schätzele, Natalie Burdack and Laurie Schiller. Special thanks to the “Nationalpark Vorpommersche Boddenlandschaft”, the “Biosphärenreservat Südost-Rügen” and the “Nationalpark Schleswig-Holsteinisches Wattenmeer” for the permissions to sample within protected areas.

This work was funded via a PhD stipend awarded to A.B. from the German Federal Environmental Foundation (DBU, AZ: 20019/635). A.B. was additionally supported via the project MV Seagrass for Climate (BMUV 67AN30003B). K.K. was supported via the project SeaStore (BMBF 03F0859C, BMUV 03F0978C, PI: M.M.B.)

