## Supplemental Table S1, S2 and S3 for "Eelgrass seeds host a distinct microbiome that is consistent along a salinity gradient"

Table S1: list of sampling locations and environmental conditions at time point of sampling

| **site** | **min_ depth [m]** | **max_ depth [m]** | **secchi_ depth [m]** | **salinity [ppm]** | **pH** | **temperature [°C]** | **Date of sampling** |
|---|---|---|---|---|---|---|---|
| Rostock | 1.8 | 2.2 | / | 10 | 7.5 | 16 | 18.06.2020 |
| Falckenstein | 0.8 | 5.0 | / | 14 | 7.9 | 16 | 23.06.2020 |
| Badeanstalt | 1.8 | 3.0 | / | 14 | 7.5 | 17 | 23.06.2020 |
| Heidkate | 1.5 | 2.0 | / | 14 | 7.5 | / | 24.06.2020 |
| Poel | 0.7 | 1.3 | / | / | / | 20.3 | 24.06.2020 |
| Schaprode1 | 2.7 | 2.7 | 1.6 | 9 | 7.9 | / | 14.07.2020 |
| Libben | 3.1 | 3.1 | >3.1 | 9.5 | 7.5 | / | 14.07.2020 |
| Having | 2.4 | 2.4 | 1.4 | 9 | / | / | 13.07.2020 |
| Vilm1 | 1.5 | 1.8 | / | 8.5 | 7.5 | / | 13.07.2020 |
| HamburgerHallig | 0.5 | / | / | 23 | 8.4 | / | 05.09.2020 |

Table S2: pairwise comparisons of bacterial community compositions between the different sample types, results rounded to two decimals

| | pairs | Df | SumsOfSqs | F.Model | R2 | p.value | p.adjusted |
|---|---|---|---|---|---|---|---|
| 1 | old leaf vs root | 1 | 1.27 | 3.49 | 0.16 | 0.001 | 0.006 |
| 2 | old leaf vs seed | 1 | 1.23 | 3.42 | 0.16 | 0.001 | 0.006 |
| 3 | old leaf vs young leaf | 1 | 0.52 | 1.39 | 0.07 | 0.077 | 0.462 |
| 4 | root vs seed | 1 | 1.48 | 4.31 | 0.19 | 0.001 | 0.006 |
| 5 | root vs young leaf | 1 | 1.40 | 3.92 | 0.18 | 0.001 | 0.006 |
| 6 | seed vs young leaf | 1 | 1.31 | 3.71 | 0.17 | 0.001 | 0.006 |

Table S3: pairwise comparisons of eukaryotic community compositions between the different sample types, results rounded to two decimals

| | pairs | Df | SumsOfSqs | F.Model | R2 | p.value | p.adjusted | sig |
|---|---|---|---|---|---|---|---|---|
| 1 | old leaf vs root | 1 | 1.17 | 2.87 | 0.14 | 1 | 6 | * |
| 2 | old leaf vs seed | 1 | 1.09 | 2.68 | 0.14 | 1 | 6 | * |
| 3 | old leaf vs young leaf | 1 | 0.38 | 0.93 | 0.05 | 551 | 1 | |
| 4 | root vs seed | 1 | 1.01 | 2.40 | 0.13 | 1 | 6 | * |
| 5 | root vs young leaf | 1 | 1.02 | 2.43 | 0.13 | 2 | 12 | . |
| 6 | seed vs young leaf | 1 | 0.96 | 2.30 | 0.14 | 2 | 12 | . |
